# Plasma metabolomics of presymptomatic *PSEN1*-H163Y mutation carriers: A pilot study

**DOI:** 10.1101/2020.05.16.093559

**Authors:** Karthick Natarajan, Abbe Ullgren, Behzad Khoshnood, Charlotte Johansson, José Miguel Laffita-Mesa, Josef Pannee, Henrik Zetterberg, Kaj Blennow, Caroline Graff

## Abstract

**Background and Objective:** *PSEN1*-H163Y carriers, at the presymptomatic stage, have reduced ^18^FDG-PET binding in the cerebrum of the brain [1]. This could imply dysfunctional energy metabolism in the brain. In this study, plasma of presymptomatic *PSEN1* mutation carriers was analyzed to understand associated metabolic changes.

**Methods:** We analyzed plasma from non-carriers (NC, n=8) and presymptomatic *PSEN1*-H163Y mutation carriers (MC, n=6) via untargeted metabolomics using gas and liquid chromatography coupled with mass spectrometry, which identified 1199 metabolites. All the metabolites were compared between MC and NC using univariate analysis, as well as correlated with the ratio of Aβ_1-42/Aβ1-40_, using Spearman’s correlation. Altered metabolites were subjected to Ingenuity Pathways Analysis (IPA).

**Results:** When comparing between presymptomatic MC and NC, the levels of 116 different metabolites were altered. Out of 116, only 23 were annotated metabolites, which include amino acids, fatty acyls, bile acids, hexoses, purine nucleosides, carboxylic acids, and glycerophosphatidylcholine species. 1-docosapentaenoyl-GPC, glucose and uric acid were correlated with the ratio of plasma Aβ_1-42_/Aβ_1-40_ (p < 0.05).

**Conclusion:** This study finds dysregulated metabolite classes, which are changed before the disease onset. Also, it provides an opportunity to compare with sporadic Alzheimer’s Disease. Observed findings in this study need to be validated in a larger and independent Familial Alzheimer’s Disease (FAD) cohort.

**Graphical abstract:** 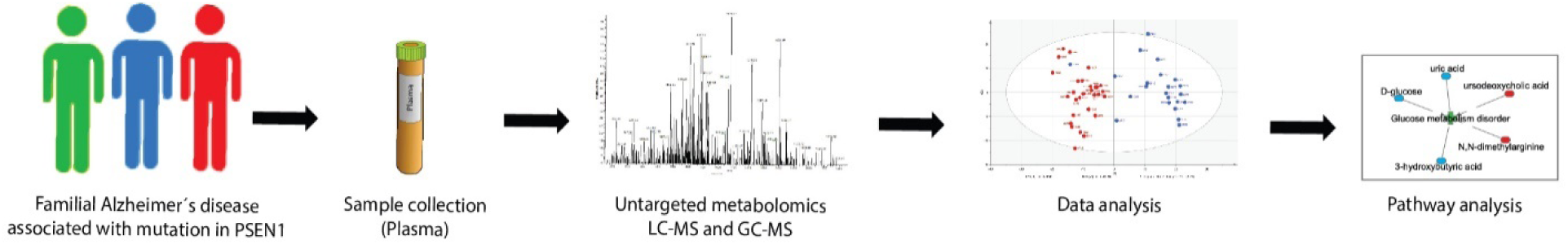

## Introduction

Alzheimer’s disease (AD) is a growing health concern estimated to affect 136 million people worldwide by 2050 [2]. The autosomal dominant mutations found in *APP, PSEN1 and PSEN2* [3] cause early-onset Familial Alzheimer’s Disease (FAD) with the high penetrance *PSEN1* mutations (https://www.alzforum.org/mutations/psen-1) being the most frequent [4]. PSEN1 is a part of the gamma-secretase protein complex, which cleaves the Amyloid precursor protein (APP) that leads to the production of a mixture of amyloid peptides Aβ_1–40_ (90%) and Aβ_1–42_ (10%) in the amyloidogenic pathway [4]. These peptides are known to be part of the pathobiology of FAD. The *PSEN*1 mutation p.His163Tyr (*PSEN1-*H163Y) results in higher levels of Aβ_1-42/_Aβ_1-40_ [5].

Previously, several studies have described the plasma metabolic profile of sporadic mild cognitive impairment (MCI) and AD cases [2, 6-12]. A targeted plasma lipid profiling was performed in *PSEN1* mutation carriers [13]. However, no untargeted plasma metabolome has yet been reported for presymptomatic *PSEN1* mutation carriers.

In FAD, the preclinical phase is characterized by brain glucose hypometabolism [14, 15]. We had previously demonstrated glucose hypometabolism in the cerebrum at the presymptomatic stage of *PSEN1*-H163Y carriers [1]. In this follow-up study, we hypothesize that untargeted plasma metabolite profiling could provide clues about the dysregulated metabolic pathways in pre-symptomatic *PSEN1* mutation carriers (MC) in comparison with non-carriers (NC) within the FAD *PSEN1*-H163Y cohort [3, 16-18]. First, we identified metabolites, which are differentially expressed between MC and NC and assessed their associated biological processes. Then, we explored the association between these metabolites with plasma levels of various amyloid beta species (Aβ). Taken together, our study provides a snapshot of the plasma metabolic changes in presymptomatic *PSEN1* MC, which may inform about biological events that are characteristics of the preclinical phase.

## Materials and Methods

### Study population

Relatives with plasma samples from the *PSEN1*-H163Y kindred in the Swedish FAD study were eligible for inclusion. The prospective FAD study invites *APP* and *PSEN1* kindreds at the Memory outpatient clinic in Karolinska University Hospital, Stockholm, since 1993. Presymptomatic relatives with 50% risk of disease are followed longitudinally with a comprehensive assessment battery, described in detail elsewhere [3, 16, 18]. Clinical and neuroradiological evaluation is accompanied by sampling of cerebrospinal fluid (CSF) and blood. Mean (SD) age of onset in the *PSEN1*-H163Y kindred is 52 ±6 years (based on 12 individuals). Symptom onset is regarded as having the first subjective symptoms as experienced by participants or next-of-kin. Mutation status (carrier or non-carrier) was not known to participants or clinicians within the study, if not stated otherwise. The study procedures were approved by the Regional Ethical Review Board in Stockholm, Sweden, and were in agreement with the Helsinki Declaration. All participants provided written informed consent to participate. The plasma samples were collected in a longitudinal manner as a part of the battery of clinical evaluation over the years from 1995 to 2017.

### DNA extraction from Blood

As part of the FAD study protocol, venous blood was collected at the Memory clinic of Karolinska university hospital, Stockholm. Using the Gentra Puregene blood Kit (Qiagen, Hilden, Germany) DNA was extracted from the blood and resuspended in RNase & DNase free water (Qiagen, Hilden, Germany). The concentration of extracted DNA was measured with the QUBIT instrument (Thermofisher, Waltham, MA, USA) as described by the manufacturer.

### Genotyping for *PSEN1*-H163Y mutation

20ng of DNA was amplified for *PSEN1*-Exon6 using forward (5’ GGTTGTGGGACCTGTTAATT 3’) and reverse (5’ CAACAAAGTACATGGCTTTAAATGA 3’) primers with AmpliTaq Gold® 360 PCR Master Mix (Thermofisher, Waltham, MA, USA). Sanger sequencing was performed using BigDye™ Terminator v3.1 Cycle Sequencing Kit (Thermofisher, Waltham, MA, USA) in both forward and reverse directions and analyzed using ABI3500 Genetic Analyzer (Thermofisher, Waltham, MA, USA).

### *APOE* allele genotyping

The *APOE* genotyping was performed for SNP rs7412 and rs429358 using pre-designed TaqMan® SNP Genotyping Assays (Thermofisher, Waltham, MA, USA) as indicated in the manufacturer’s protocol. The amplified products were run on 7500 fast Real-Time PCR Systems (Thermofisher, Waltham, MA, USA).

### Plasma sample collection

Non-fasting plasma was prepared from the blood. The plasma was removed as the supernatant after 1hr incubation at room temperature (RT) and 10 mins centrifugation at 2200g. Then, the plasma was aliquoted and frozen at −80°C until analysis. Plasma sampled between the years 1995 to 2017 were included. The metabolite analysis was carried out at the Swedish Metabolomics Center (SMC, https://www.swedishmetabolomicscentre.se/), Umeå, Sweden.

### Metabolite extraction and analysis

Untargeted metabolite extraction and analysis via Gas Chromatography (GC) and Liquid Chromatography (LC) in combination with Mass-Spectrometry (MS) were performed at SMC as described here [19]. Plasma was prepared by adding 900µl of extraction buffer (90/10 v/v methanol: water) along with internal standards for the GC-MS and LC-MS to 100µl of plasma. The samples were prepared and analyzed in a randomized order for GC-MS and LC-MS. Detailed methods were described in the supplementary methods section (Metabolite profiling of the plasma).

### Analysis of Aβ_1-38_, Aβ_1-40_, and Aβ_1-42_ in plasma

The analyses of Aβ_1-38_, Aβ_1-40_, and Aβ_1-42_ in plasma were performed using immunoprecipitation coupled to tandem Liquid Chromatography mass spectrometry (IP-LC-MS/MS) as described previously [20, 21]. In short, calibrators were prepared using recombinant Aβ1-38, Aβ1-40 and Aβ1-42 (rPeptide) added to 8 % bovine serum albumin in phosphate-buffered saline. Recombinant ^15^N uniformly labeled Aβ_1-38_, Aβ_1-40_ and Aβ_1-42_ (rPeptide) were used as internal standards (IS), added to samples and calibrators prior to sample preparation. Aβ peptides were extracted from 250 µL human plasma using immunoprecipitation with anti-β-Amyloid 17-24 (4G8) and anti-β-Amyloid 1-16 antibodies (6E10, both Biolegend®) coupled to Dynabeads™ M-280 Sheep Anti-Mouse IgG magnetic beads (Thermofisher, Waltham, MA, USA). Immunoprecipitation was performed using a KingFisher™ Flex Purification System (Thermofisher, Waltham, MA, USA).

Analysis of processed samples was performed using liquid chromatography-tandem mass spectrometry (LC-MS/MS) on a Dionex Ultimate LC-system and a Thermo Scientific Q Exactive quadrupole-Orbitrap hybrid mass spectrometer. Chromatographic separation was achieved using basic mobile phases and a reversed-phase monolith column at a flow rate of 0.3 mL/min. The mass spectrometer operated in parallel reaction monitoring (PRM) mode was set to isolate the 4+ charge state precursors of the Aβ peptides. Product ions (14-15 depending on peptide) specific for each precursor was selected and summed to calculate the chromatographic areas for each peptide and its corresponding IS. The area ratio of the analyte to the internal standard in unknown samples and calibrators was used for quantification.

### Statistical analysis

All statistical analyses were done using R (The R Foundation for Statistical Computing; version 3.6.1) and R Studio software. Group comparisons between presymptomatic *PSEN1*-H163Y MC and NC were made using Wilcoxon rank-sum tests and a p value below 0.05 was considered as significant. Features with p values above the significance threshold were excluded from downstream analysis. Correlations between the selected metabolites and plasma Aβ_1-42/_Aβ_1-40_ ratio were tested using Spearman’s rho statistic and the p values were calculated using the asymptotic t approximation.

The correlation between the observed metabolite levels and how many years the sample had been kept in storage was tested using Spearman’s rho statistic and the p values were calculated using the asymptotic t approximation. The same was done for correlations with participant age at sampling and the presence of at least one *APOEε4* allele. The heatmaps were constructed using two separate clusters, one cluster for the features and another for the samples. The clustering analysis was done using agglomerative hierarchical clustering using the Wards clustering criterion. The dissimilarity matrices were constructed using Pearson’s correlation.

### Metabolites biological interpretation

Differential metabolites between MC and NC were converted into their corresponding Human Metabolome Database identifier (http://www.hmdb.ca/, HMDB ID) as well as annotated for their metabolite class. HMDB ID was analyzed using the Ingenuity Pathway Analysis (IPA) (https://www.qiagenbioinformatics.com/products/ingenuity-pathway-analysis/, Qiagen Inc.) software, enriched significant metabolite ontologies were filtered as described in the IPA’s metabolomics white paper (http://pages.ingenuity.com/rs/ingenuity/images/wp_ingenuity_metabolomics.pdf).

## Results

### Demographics of the study population

A total of 24 plasma samples from 17 males were included in this study, of these there were 6 mutation carriers (MC) and 11 non-carriers (NC). The study participants were relatives from the *PSEN1*-H163Y kindred, except for three NC from two *APP* kindreds (Table 1). The participants underwent repeated sampling of blood, meaning these individuals were represented more than once and contributing to an overall mean age at sampling of 42 ± 11 years. Distribution of mutation status, *APOEε4* status and mean age at sampling in each dataset is described in table 1. Asymptomatic status in all participants was confirmed by later records of the actual age of onset in the MC and mean Mini-Mental State Examination (MMSE) scores [22] were 29 ±1 (maximum score 30) upon sampling. One of the participants from the MC group opted for presymptomatic genetic testing at the hospital. This has since proved to be a rare case of reduced penetrance, showing no signs of beta-amyloid (Aβ) retention during [11C]Pittsburgh compound B (PiB) positron emission tomography (PET) at the age of 60 years [16] and no cognitive deficits during the most recent psychological assessment, performed at the age of 65, 13 years past the expected age of onset. There were no known subjects with alcohol overconsumption or dietary restrictions.

**Table 1.**
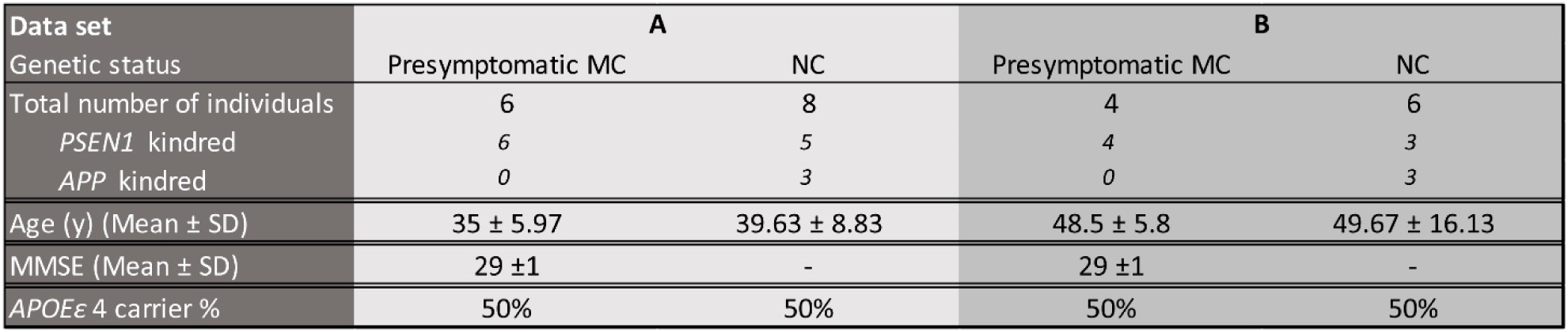
Demographic data for dataset A (samples collected before 2008) and dataset B (samples collected after 2008), which were used for the univariate analysis. Plasma samples were stratified for univariate analysis based on the PCA (Supplementary fig.1). One “baseline” plasma sample from each time period was selected, and all individuals were male and presymptomatic upon sampling. The number of Apolipoprotein E4 (*APOEε4*) genotype carriers in each dataset is indicated. Seven of the participants (3 NC and 4 MC) are represented in both A and B datasets (‘y’ denotes years).

### Comparison of plasma metabolite levels between presymptomatic MC and NC

Untargeted GC-MS and LC-MS analysis of the plasma samples detected 1199 metabolites, of which 23% were annotated. First, a Principal Component Analysis (PCA) model was built (Supplementary fig.1). The PCA analysis indicate a clear separation of the samples based on the number of years of storage. A similar effect was previously reported [23]. For that reason, the samples were stratified into two separate data sets based on the sample collection year (Table 1; referred to as data set A and B in the text). Dataset “A” contains the samples collected before the year 2008 and dataset “B” contains the samples collected after 2008.

The two data sets were analyzed separately. Within them the measured levels of the metabolites were compared between the MC and NC. In dataset A, the levels of 79 metabolites (14 annotated), were significantly different in MC compared with NC (P < 0.05) (Table 2, Supplementary table 1). The annotated metabolites belong to the metabolite classes of amino acids (n = 3), carboxylic acids (n = 1), hexoses (n = 1), imidazopyrimidines (n = 1), fatty acyls (n = 1), glycerophospholipids (n = 6) and hydroxy acids (n = 1). In dataset B, the levels of 37 metabolites (9 annotated), were significantly different in MC compared with NC (P < 0.05) (Table 2, Supplementary table 1). The annotated metabolites belong to the metabolite classes of amino acids (n = 2), purine nucleosides (n = 1), carboxylic acids (n = 2), fatty acyls (n = 2) and bile acids (n = 2).

**Table 2.** Annotated metabolites (n=23) showed a significant (p<0.05) difference between MC and NC. Log2 foldchange is shown as the difference between MC and NC, with positive values indicating higher relative metabolite levels in MC, and vice versa. § The metabolite is present in both dataset A and dataset B. *Significant correlation (p<0.05) between metabolite and Aβ peptides.

| Data set | Metabolite | Metabolite class | Fold change | p.value | Correlation to Aβ(1-42/1-40) |
| --- | --- | --- | --- | --- | --- |
| B | Glycohyocholic acid | Bile acids | -1.023 | 0.014 | -0.236 |
| B | Asymmetric dimethylarginine | Carboxylic acids | 0.403 | 0.025 | 0.418 |
| B | Ursodeoxycholic acid | Bile acids | 2.193 | 0.025 | -0.006 |
| B | Threonine § | Amino acids | 0.392 | 0.043 | 0.042 |
| B | Asparagine | Amino acids | 0.417 | 0.043 | 0.164 |
| B | N2,N2-Dimethylguanosine | Purine nucleosides | 0.649 | 0.043 | 0.139 |
| B | N-Acetylvaline | Carboxylic acids | 0.341 | 0.043 | -0.018 |
| B | 3-Methylglutaryl carnitine | Fatty acyls | -0.793 | 0.043 | -0.382 |
| B | 2-Hydroxycaproic acid | Fatty Acyls | 0.339 | 0.043 | 0.394 |
| A | 1-oleoyl-GPC | Glycerophospholipids | 0.170 | 0.003 | 0.544 |
| A | Cystine | Amino acids | -0.289 | 0.017 | -0.341 |
| A | Glutamine | Amino acids | 0.157 | 0.017 | 0.505 |
| A | 2-oleoyl-GPC | Glycerophospholipids | 0.335 | 0.017 | 0.505 |
| A | Pyroglutamic acid | Carboxylic acids | 0.153 | 0.024 | 0.544 |
| A | Uric acid | Imidazopyrimidines | -0.496 | 0.024 | -0.609* |
| A | 1-Palmitoyl-sn-glycero-3-phosphocholine | Glycerophospholipids | 0.436 | 0.024 | 0.390 |
| A | 1-arachidonoyl-GPC | Glycerophospholipids | 0.337 | 0.024 | 0.401 |
| A | 3-Hydroxybutyric acid | Hydroxy acids | -0.731 | 0.024 | -0.341 |
| A | Glucose | Hexoses | -0.164 | 0.033 | -0.571* |
| A | Threonine § | Amino acids | 0.138 | 0.045 | 0.445 |
| A | Propionyl carnitine | Fatty Acyls | -0.497 | 0.045 | -0.374 |
| A | 2-stearoyl-GPC | Glycerophospholipids | 0.264 | 0.045 | 0.220 |
| A | 1-docosapentaenoyl-GPC | Glycerophospholipids | 0.342 | 0.045 | 0.648* |

In addition, the correlation between all the metabolites that showed a significant difference between NC and MC in the two data sets, and the number of years that the samples have been in storage was tested. Years in storage was calculated as the difference between sampling date and analysis date. This was done to investigate if the observed differences could be due to the above described storage effect (Supplementary table 2). In dataset A, one unannotated metabolite had a significant correlation with years in storage. This metabolite was excluded from the cluster analysis and the pathway analysis. In dataset B, no metabolites had a significant association with years in storage. Similar correlation analyses were done between metabolite levels and participant age at the time of sampling, as well as the presence of at least one allele of *APOEε4* (Supplementary table 2). In dataset A, pyroglutamic acid and 2-oleoyl-GPC had a significant correlation with age at sampling (p=0.027 and p=0.043, respectively). In dataset B, no significant correlation was observed between any of the annotated metabolites and age at sampling. No significant correlation of metabolites with the presence of *APOEε4* was observed in either set.

Heatmaps (Fig.1A-B) for the two datasets were generated based on the hierarchal clustering of both the samples and the metabolites (Table 2, Supplementary table 1). In both datasets, the MC and the NC separate into two different clusters. Furthermore, the metabolites form two distinct clusters in both datasets as well, where one cluster of metabolites was found at higher levels in MC than in NC and the other shows the opposite trend (Fig.1A-B).

**Fig.1:**
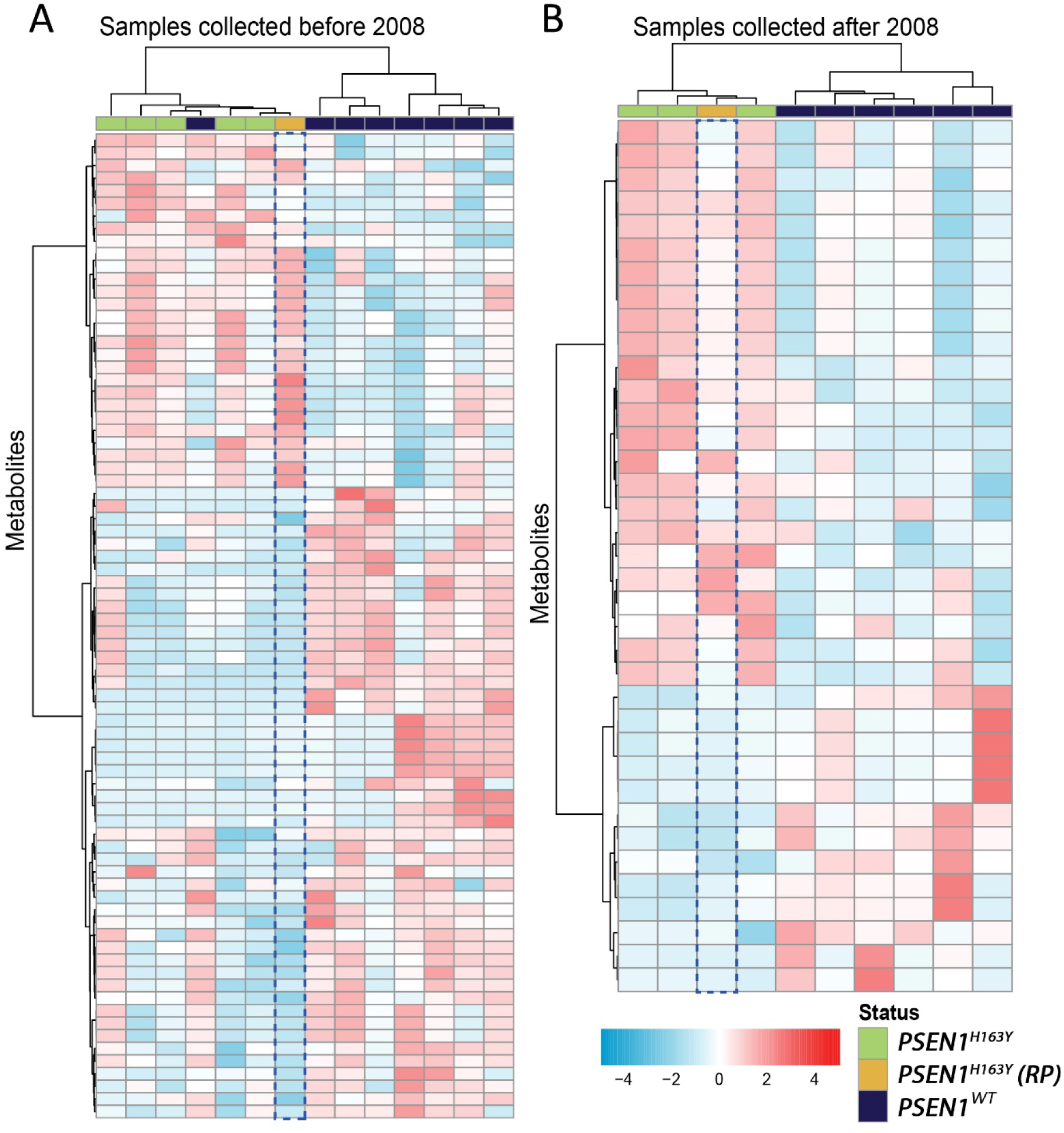
Heat map showing similarities between samples of the metabolites that were differentially expressed between NC (non-carrier group, dark blue) and the *PSEN1-*H163Y MC (mutation carrier group, Green). Sample from the reduced penetrance mutation carrier (RP) case is represented by yellow color. Legend: Red (upregulated metabolite), Blue (downregulated metabolite) and white (no modulation). A) Samples from dataset A. B) Samples from dataset B.

### Correlations between plasma metabolites and plasma Aβ_1-42/_Aβ_1-40_ ratio

Amyloid pathology is a hallmark of AD and Aβ levels in plasma are used as possible surrogate biomarkers [24, 25]. It has been shown that *PSEN1*-H163Y exhibit higher plasma Aβ_1-42_/Aβ_1-40_ levels [5]. We, therefore, tested the correlation between all 1199 metabolites and the ratio of Aβ_1-42/_Aβ_1-40_ in plasma (Supplementary table 3). In dataset A, significant correlations were observed between 50 metabolites (12 annotated) and the ratio of Aβ_1-42/_Aβ_1-40_ (Table 3). In dataset B, significant correlations were observed between 55 metabolites (18 annotated) and the ratio of Aβ1_-42/_Aβ_1-40_ (Table 3). This includes glucose, uric acid and 1-docosapentaenoyl-GPC, which exhibit differential levels in MC compared to NC (Table 3).

**Table 3.**
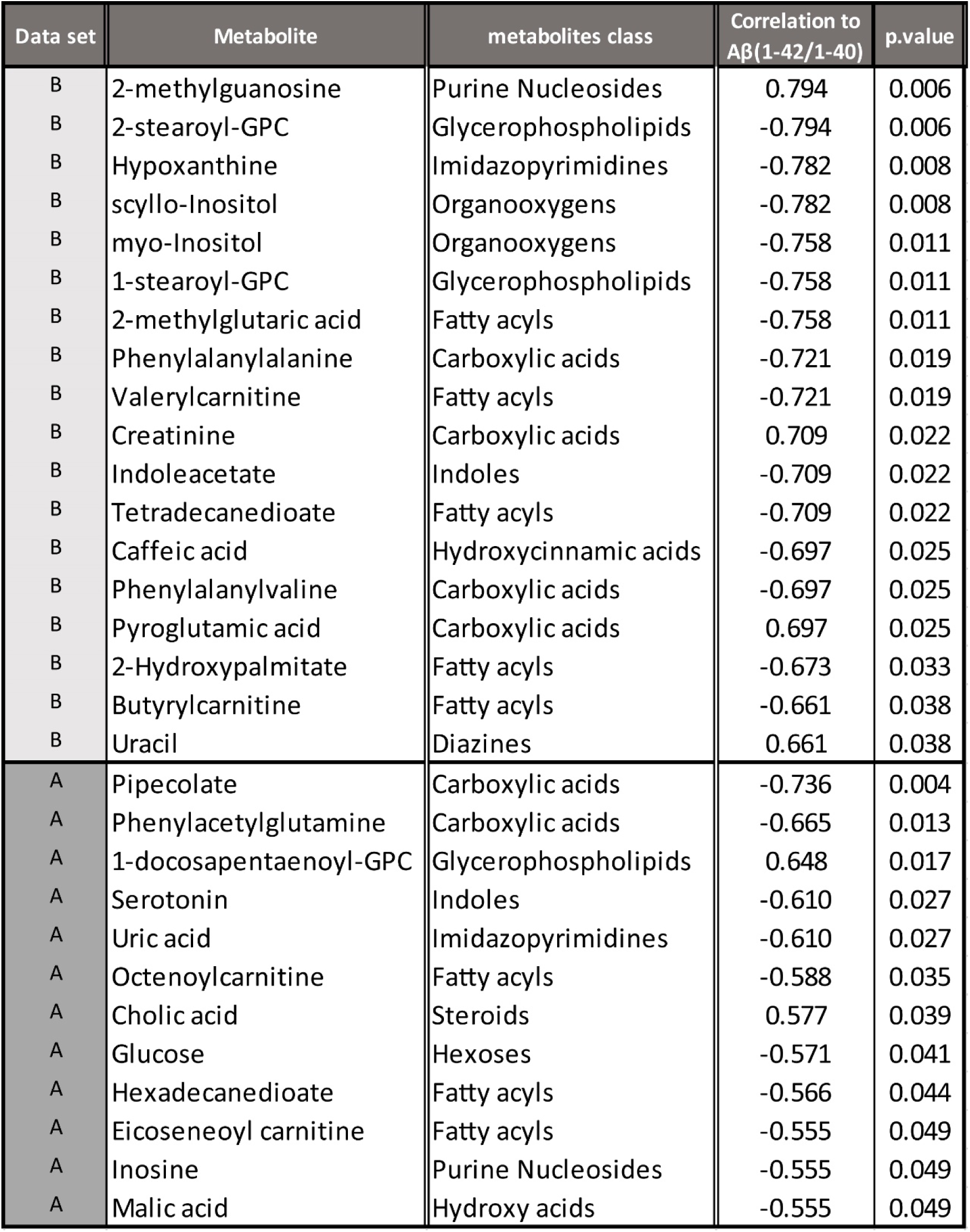
Spearman correlation between 1199 detected metabolites, in both dataset A and B, and Aβ_1-42/1-40_-ratio. The table shows the annotated metabolites that show a significant correlation (p<0.05).]

### Biological significance of annotated metabolites

To examine the biological significance of 23 annotated metabolites from dataset A and dataset B, we performed the Ingenuity Pathway Analysis (IPA). Canonical pathways identified include ‘Asparagine Biosynthesis I’, ‘tRNA Charging’, ‘Asparagine Degradation I’, ‘Glutamine Degradation I’ and ‘IL-12 signaling and production in Macrophages’ (Fig.2A). Along with pathway enrichment, the IPA also enriched metabolites for two different categories ‘Disease and Disorders’ and ‘Molecular and Cellular Functions’ (Supplementary fig. S1). We explored these categories to understand the role of these metabolites in the presymptomatic MC. In ‘Molecular and Cellular functions’, the metabolites were part of the cellular process ‘Lipid metabolism’, ‘Cellular Growth and proliferation’ (Supplementary fig.S1), ‘Peroxidation of lipid’, ‘Production of reactive oxygen species’, and ‘Neuronal cell death’ (Fig.2B-D). In addition, under the category ‘Disease and Disorders’ the metabolites were linked to the disease-related phenotypes including ‘Metabolic disease’, ‘Endocrine system disorder’, and ‘Chronic inflammatory disorder’ (Fig.2E-F, Supplementary fig.S1).

**Fig.2:**
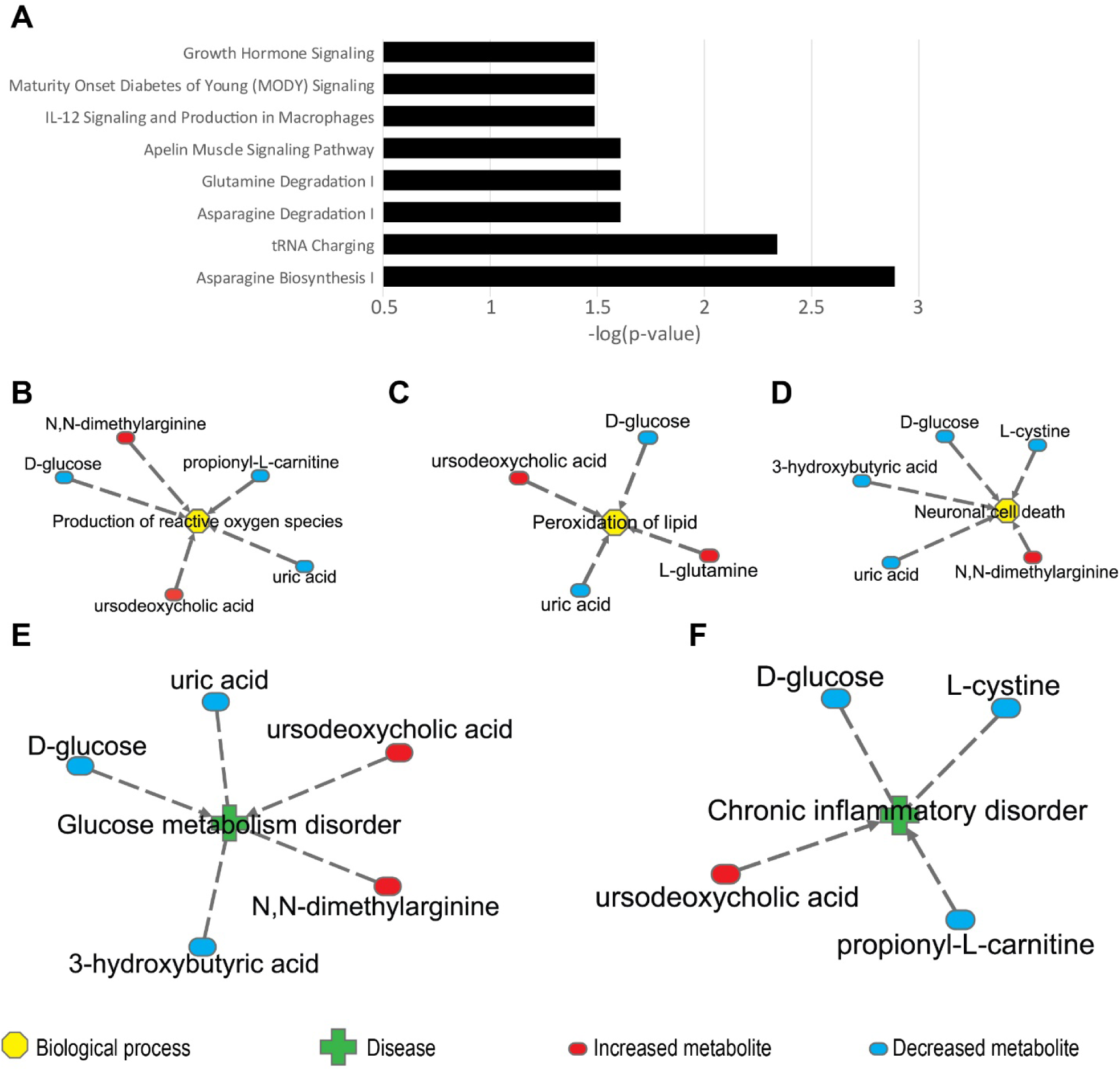
A) Enriched canonical pathways for 23 annotated metabolites, which were differentially expressed between presymptomatic *PSEN1* MC and NC. B-D). In ‘Molecular and Cellular Functions’ terms, identified metabolites were associated with B) ‘Production of reactive oxygen species’ C) ‘Peroxidation of lipid’, D) ‘Neuronal cell death’. E-F). In ‘Disease and Disorders’ terms, under ‘Metabolic disease’ E) ‘Glucose metabolism disorder’ and in ‘Inflammatory disease’ F) ‘Chronic inflammatory disorder’ metabolite network was identified.

Metabolites are the functional endpoint of biological processes, which do not act alone and are regulated by upstream regulators at a given physiological condition [26, 27]. Therefore, with the help of the ‘Upstream Regulator Analysis’ algorithm part of IPA, (supported by the Ingenuity® Knowledge Base) various, statistically significant, regulators of the 23 metabolites were identified (Supplementary table S4). These upstream regulators include the transcription factors TFE3 (Transcription factor E3), ZBTB20 (Zinc finger- and BTB domain-containing protein 20) and CEBPB (CCAAT-Enhancer Binding Protein-β). Moreover, along with transcription factors, IPA identified the enzymes DDAH2 (dimethylarginine dimethylaminohydrolase-2), FMO3 (flavin monooxygenase-3) and PIK3CA (phosphatidylinositol-4,5-bisphosphate 3-kinase alpha), which are part of regulatory networks. Further, metabolites that are part of ‘Lipid metabolism’ and ‘Cell cycle’ were visualized using IPA (Fig.3A-B).

**Fig.3:**
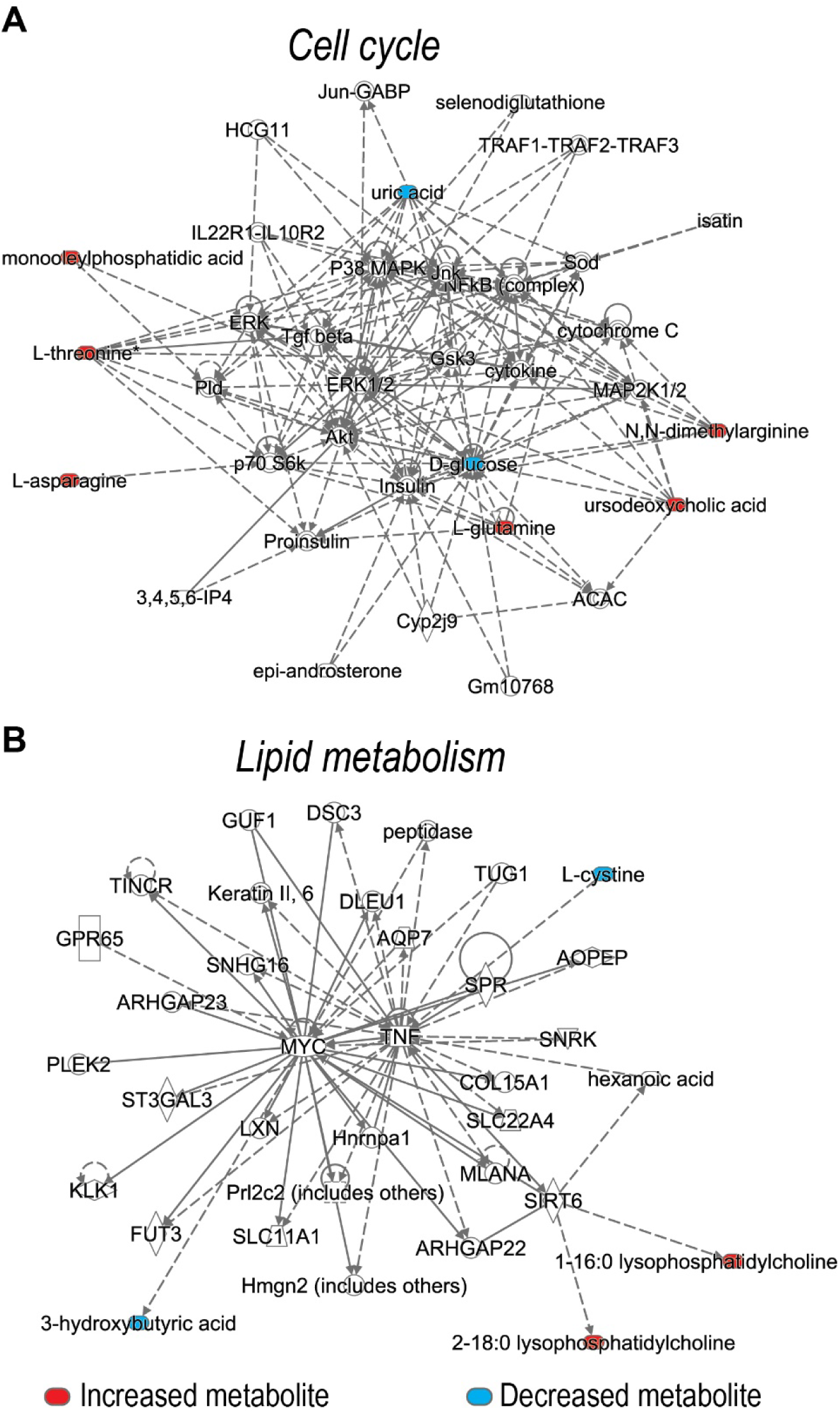
A) Network of metabolites and its regulators involved in the ‘Cell cycle’. B) Molecular network of differentially expressed metabolites and their upstream regulators involved in the ‘Lipid metabolism’.

## Discussion

In this study, we performed an untargeted screening of plasma metabolites in male presymptomatic *PSEN1-*H163Y MC known to exhibit brain glucose hypometabolism early in the preclinical phase, up to 20 years prior to expected onset age [1]. *APOEε4* is represented in both MC and NC and metabolite correlation analysis did not indicate skewness [28]. In presymptomatic MC, amino acids, fatty acyls, carboxylic acids, hexoses, purine nucleosides, glycerophosphatidylcholines, and bile acids were significantly altered when compared to NC (Table 2). Interestingly, these classes of metabolites are also associated with sporadic AD [2, 6, 8, 9, 11, 12, 29-32]. Notably, we found asparagine, propionyl-l-carnitine, glycerophosphatidylcholine species and asymmetric dimethylarginine (Table 2), which were part of the metabolite panel that previously has been reported to predict the clinical transition from presymptomatic to prodromal or symptomatic late-onset AD[9].

We identified differential metabolites associated with ‘Production of reactive oxygen species’, ‘lipid peroxidation’, ‘glucose metabolism disorder’ and ‘chronic inflammatory disorder’ (Fig.2, Supplementary fig.S1) which are known to be inherent of FAD pathobiology [15, 33-36]. Amyloid-beta mediated mitochondrial dysfunction associated with glucose hypometabolism is an indicator of molecular events, which precede amyloid aggregation [37-39]. Oligomeric forms of Aβ can give rise to oxidative stress placing themselves in the lipid bilayer which causes lipid peroxidation, at the end leading to neuronal cell death [15, 40, 41]. Furthermore, increased levels of 1-docosapentaenoyl-GPC, decreased levels of glucose and uric acid in MC were significantly correlated with Aβ_1-42_/Aβ_1-40_ (Table 2). Besides, several metabolites were significantly correlated with Aβ_1-42/Aβ1-40,_ but not identified in the univariate analysis, probably due to a lack of statistical power (Supplementary table 3). The suggested upstream regulators of the identified metabolites in our study are proposed to play a role in modulating different cellular events. In the event of mitochondrial dysfunction, TFE3 is activated and plays a vital role in regulating energy metabolism [42]. In addition, ZBTB20 is another upstream regulator that plays an essential role in glucose homeostasis [43].

In the context of inflammation, astrocytes and microglia are the principal players in AD [26, 44] and their activation is characteristic of the preclinical phase [45]. Aβ activates microglia (Brain resident macrophages) which trigger a proinflammatory cascade, likely IL-12, and IL-23 [46, 47]. The metabolites found in our study (Fig.2A) are associated with the IL-12 pathway. Activated microglia induce neurotoxic reactive astrocytes [48]. Moreover, the metabolite upstream regulator CEBPB identified here, is activated by Aβ in glial cells, which in turn initiates the inflammatory cascade [49].

A rare case of *PSEN1*-H163Y reduced penetrance (RP) [16] is also part of our study. We included the plasma samples collected before the age of onset (52±6) and the RP case exhibits a similar metabolic profile to the other MC (Fig.1). Such reduced penetrance cases are rare and have been shown to have other protective genetic or environmental modifiers [50]. Further experimental studies employing single-cell genomics [51] and directly induced neurons from fibroblasts [52] can shed more light on possible mechanisms leading to resilience.

This pilot-scale study has its limitations. The study design is observational, it is difficult to control all the confounding factors, for example, the metabolite profile is highly influenced by lifetime immunological experience [53], gut microbiota [54] and exposome [55]. Also, the medication history of the subjects was not accounted for. Considering the low frequency of the *PSEN1* mutation carriers in our Swedish cohorts the sample size was small, and the p values were not corrected for multiple comparisons in the univariate analysis. It has been shown that blood metabolites are strongly associated with age, yet their association is highly selective [56]. In our analysis, we found pyroglutamic acid and 2-oleoyl-GPC to be correlated with age (Supplementary table 2).

Here, we report plasma metabolites, their upstream regulators and pathways, which were dysregulated in presymptomatic *PSEN1*-H163Y mutation carriers. These are consistent with previous findings of FAD pathophysiology. In this study, we found several unannotated metabolites [56] (Supplementary table 1 & table 3) that will be of significant interest to future FAD metabolomics studies. However, results from this pilot study need to be evaluated in a larger FAD cohort. Considering the low frequency of FAD cases in the Swedish population, acquiring the optimal sample size for similar studies remains the biggest hurdle. The FAD cases can be highly informative on the cellular and molecular front when metabolomics are integrated with multi-omics datasets [27, 57].

## Data availability

Datasets can be accessed at MetaboLights (https://www.ebi.ac.uk/metabolights/) with identifier MTBLS1721.

## Acknowledgments

Swedish Metabolomics Centre, Umeå, Sweden (www.swedishmetabolomicscentre.se) is acknowledged for metabolic profiling by GC-MS and LC-MS. The study was supported by grants provided by Swedish Alzheimer’s foundation, Brain Foundation, Demensfonden, Karolinska geriatric foundation, Stiftelsen för Gamla Tjänarinnor, Stohnes foundation, Swedish research council (2015-02926 and 2018-02754), Karolinska Institutet StratNeuro postdoc funding, and ALF Region Stockholm (County Council), Schörling Foundation - Swedish FTD Initiative.

## Authors contribution

KN, CG conceptualized and designed the study. AU and BK performed data analysis. JL performed genotyping analysis. CJ contributed to clinical data collection and evaluation. JP, HZ, and KB contributed to plasma analysis of amyloid beta. All the authors contributed to the writing of the manuscript.

## Tables, Figures & legends

**Supplementary Fig.1:**
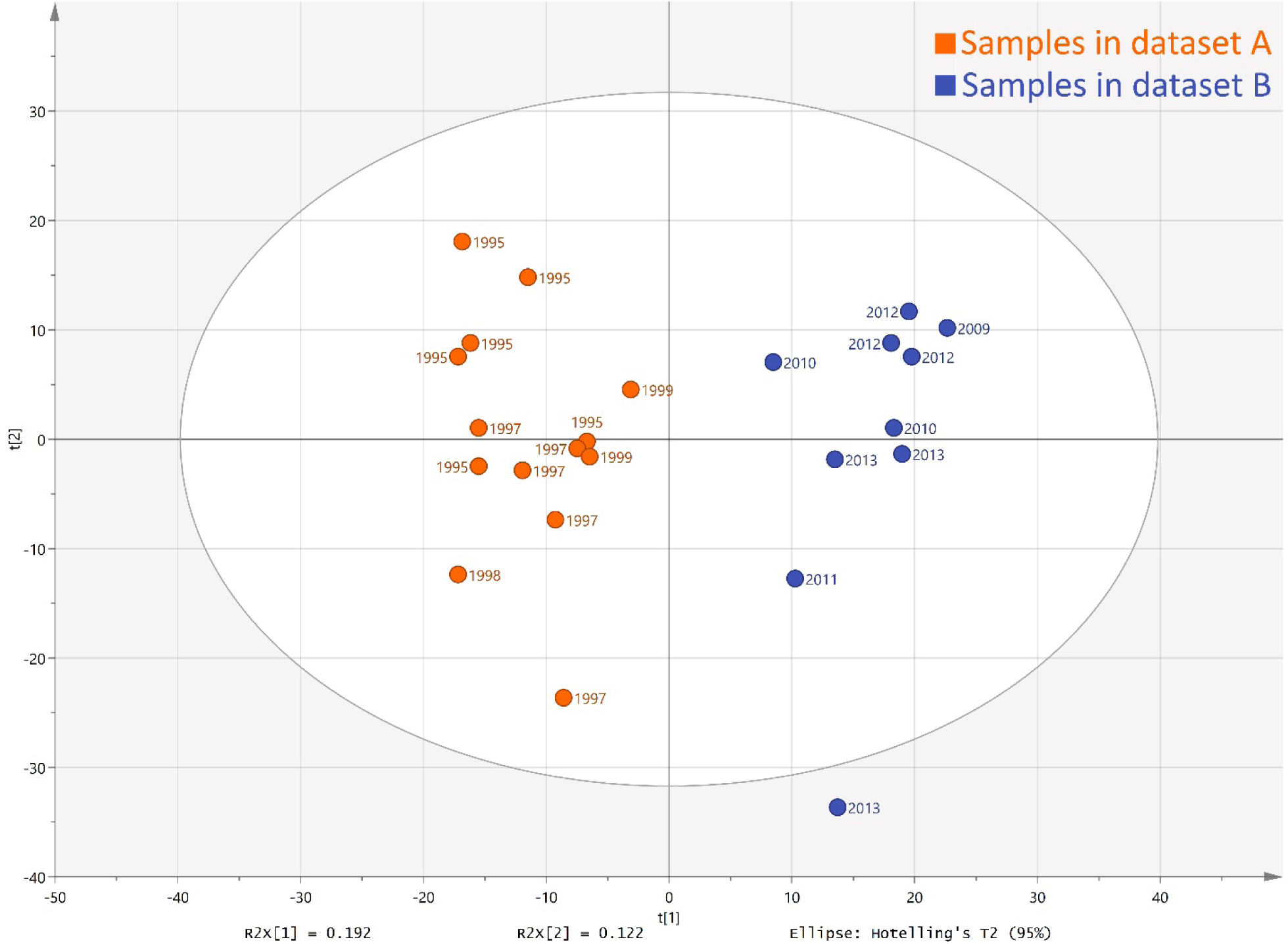
PCA of all 24 plasma samples in the study, which were analyzed for 1199 metabolites in an untargeted manner. The plasma samples used in this study were collected from the year 1995 to 2017.

**Supplementary Fig.2.**
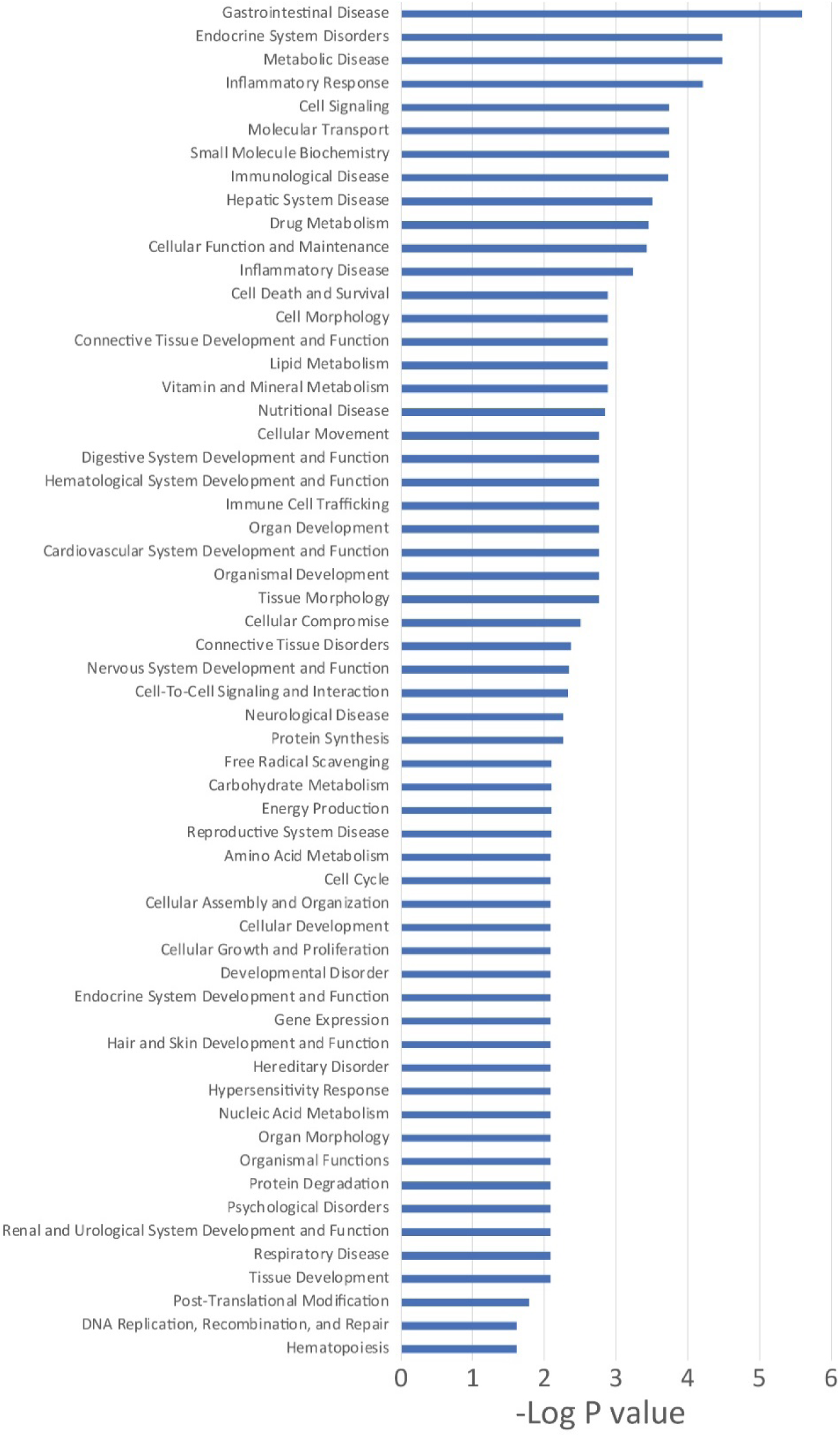
List of GO terms under the category of ‘Molecular and Cellular Functions’ and ‘Disease and Disorders’, which were enriched for the 23 annotated metabolites in the Ingenuity Pathway Analysis.

**Supplementary Table 1:** The unannotated metabolites presented as ‘mass@retention time’, that showed a significant (p<0.05) difference between MC and NC. Log2 foldchange shown as the difference between MC and NC, with positive values indicating higher relative metabolite levels in MC, and vice versa. * Significant correlation (p<0.05) between metabolite and Aβ peptides.

| Data set | Metabolite<br>(Mass@retention) | Fold<br>change | p.value | Correlation to<br>Aβ(1-42/1-40) | Data set | Metabolite<br>(Mass@retention) | Fold<br>change | p.value | Correlation to<br>Aβ(1-42/1-40) |
| --- | --- | --- | --- | --- | --- | --- | --- | --- | --- |
| B | 251.1521@6.24 | 0.571 | 0.014 | 0.382 | A | 227.0247@3.42 | -0.880 | 0.012 | -0.445 |
| B | 285.1369@5.18 | -1.048 | 0.014 | -0.6 | A | 658.2827@3.94 | -1.857 | 0.015 | -0.430 |
| B | 1240.2028@5 | 0.586 | 0.014 | 0.418 | A | 1062.6665@6.13 | 0.410 | 0.017 | 0.516 |
| B | 1239.7719@5 | 0.616 | 0.014 | 0.418 | A | 278.1885@6.09 | 0.598 | 0.017 | 0.522 |
| B | 2893.4679@5.01 § | 0.540 | 0.014 | 0.406 | A | 550.2959@5.89 | -3.478 | 0.021 | -0.435 |
| B | 2893.1341@5.01 § | 0.490 | 0.014 | 0.479 | A | 312.2121@5.52 | -0.853 | 0.024 | -0.454 |
| B | 2242.8802@4.98 | -0.096 | 0.014 | -0.467 | A | 366.2765@6.55 | -0.764 | 0.024 | -0.049 |
| B | 370.2192@2.95 § | -0.227 | 0.014 | -0.430 | A | 230.0765@1.19 | -1.215 | 0.024 | -0.341 |
| B | 356.2706@5.98 | 1.558 | 0.021 | 0.081 | A | 521.3487@6.39 | 0.485 | 0.024 | 0.440 |
| B | 392.2923@5.97 | 2.207 | 0.025 | -0.018 | A | 481.3535@6.54 | 0.414 | 0.024 | 0.390 |
| B | 1239.9154@5 | 0.554 | 0.025 | 0.261 | A | 414.2453@3.06 | -1.472 | 0.024 | -0.341 |
| B | 1240.0582@5 | 0.569 | 0.025 | 0.261 | A | 458.2716@3.16 | -1.689 | 0.024 | -0.302 |
| B | 1240.3454@5 | 0.599 | 0.025 | 0.236 | A | 470.3598@6.91 | -1.788 | 0.024 | -0.363 |
| B | 285.1363@5.28 | -0.862 | 0.025 | -0.576 | A | 482.3593@6.85 | -1.236 | 0.024 | -0.264 |
| B | 2242.3803@4.98 | -0.156 | 0.025 | -0.552 | A | 634.3765@3.43 | -0.549 | 0.029 | -0.463 |
| B | 2183.851@4.97 | 0.649 | 0.025 | 0.539 | A | 590.3503@3.37 | -1.542 | 0.029 | -0.384 |
| B | 166.0489@1.61 | 1.759 | 0.043 | -0.006 | A | 496.2309@4.02 | -5.104 | 0.029 | -0.316* |
| B | 463.1833@4.05§ | -0.787 | 0.043 | -0.672* | A | 235.0156@2.64 | -2.043 | 0.031 | -0.348 |
| B | 344.078@2.53 | 0.669 | 0.043 | 0.188 | A | 340.2385@6.85 | 1.084 | 0.032 | 0.287 |
| B | 332.0675@2.17 | 0.251 | 0.043 | 0.164 | A | 137.0461@0.4 | -1.926 | 0.033 | -0.465 |
| B | 2169.6025@5 | 0.684 | 0.043 | 0.236 | A | 458.3595@6.8 | -0.903 | 0.033 | -0.225 |
| B | 2170.3561@5 | 0.740 | 0.043 | 0.236 | A | 190.0467@1.08 | -0.339 | 0.033 | -0.148 |
| B | 1281.3591@4.99 | -0.318 | 0.043 | -0.127 | A | 543.3329@6.12 | 0.362 | 0.033 | 0.379 |
| B | 2242.6281@4.98 | -0.257 | 0.043 | -0.6 | A | 1266.4967@4.95 | 0.460 | 0.033 | 0.648* |
| B | 2242.1291@4.98 | -0.183 | 0.043 | -0.491 | A | 370.2192@2.95 § | -1.371 | 0.033 | -0.418 |
| B | 326.1933@2.79 § | -0.202 | 0.043 | -0.297 | A | 502.2981@3.24 | -1.708 | 0.033 | -0.357 |
| B | 2184.6055@4.97 § | 0.625 | 0.043 | 0.430 | A | 179.0582@2.63 | -3.790 | 0.033 | -0.34* |
| B | 463.1834@4.05 | -0.804 | 0.043 | -0.66* | A | 104.0473@1.19 | -0.705 | 0.033 | -0.313* |
| A | 194.1154@1.81 | -0.433 | 0.004 | -0.346 | A | 484.3755@6.78 | -0.732 | 0.033 | 0.033 |
| A | 2184.6055@4.97 § | 0.506 | 0.004 | 0.659* | A | 202.0298@3.69 | -1.135 | 0.033 | -0.225 |
| A | 622.3357@5.76 | -1.940 | 0.005 | -0.496 | A | 464.3494@7.19 | -0.402 | 0.033 | 0.077 |
| A | 392.2924@6.84 | -0.939 | 0.006 | -0.286 | A | 346.1526@4.39 | -0.726 | 0.037 | -0.277 |
| A | 622.3344@5.76 | -3.462 | 0.007 | -0.459 | A | 218.0248@3.54 | -1.433 | 0.043 | -0.318 |
| A | 366.2763@6.65 | -0.473 | 0.008 | -0.374 | A | 172.014@2.64 | -1.556 | 0.043 | -0.404 |
| A | 507.3688@6.63 | 0.436 | 0.008 | 0.412 | A | 135.0682@2.63 | -3.276 | 0.044 | -0.326 |
| A | 1100.6119@6.12 | 0.258 | 0.008 | 0.576* | A | 463.1833@4.05 § | -0.707 | 0.045 | -0.149 |
| A | 440.3496@7.04 | -0.956 | 0.008 | -0.346 | A | 85.089@0.62 | 0.546 | 0.045 | 0.223 |
| A | 416.2051@4.47 | -0.804 | 0.008 | -0.374 | A | 528.4017@7.19 | -0.662 | 0.045 | -0.159 |
| A | 314.2453@6.85 | -0.670 | 0.008 | -0.379 | A | 523.3646@6.82 | 0.142 | 0.045 | 0.275 |
| A | 510.2458@4.12 | -1.558 | 0.010 | -0.424 | A | 569.3488@6.21 | 0.344 | 0.045 | 0.637* |
| A | 521.3489@6.46 | 0.211 | 0.012 | 0.554* | A | 525.2866@6.1 | -0.418 | 0.045 | -0.407 |
| A | 1016.6811@6.38 | 0.418 | 0.012 | 0.533 | A | 217.1313@1.37 | -0.501 | 0.045 | -0.374 |
| A | 166.0157@0.9 | 0.140 | 0.012 | 0.214 | A | 1062.6643@6.05 | 0.651 | 0.045 | 0.418 |
| A | 2893.4679@5.01 § | 0.265 | 0.012 | 0.473 | A | 1012.6492@6.05 | 0.311 | 0.045 | 0.500 |
| A | 2893.1341@5.01 § | 0.327 | 0.012 | 0.385 | A | 326.1933@2.79 § | -1.047 | 0.045 | -0.297 |
| A | 136.0522@2.2 | -0.617 | 0.012 | -0.187 | A | 116.0472@1.43 | -0.435 | 0.045 | -0.099 |
| A | 204.0092@2.69 | -3.171 | 0.012 | -0.352 |  |  |  |  |  |

**Supplementary Table 2:**
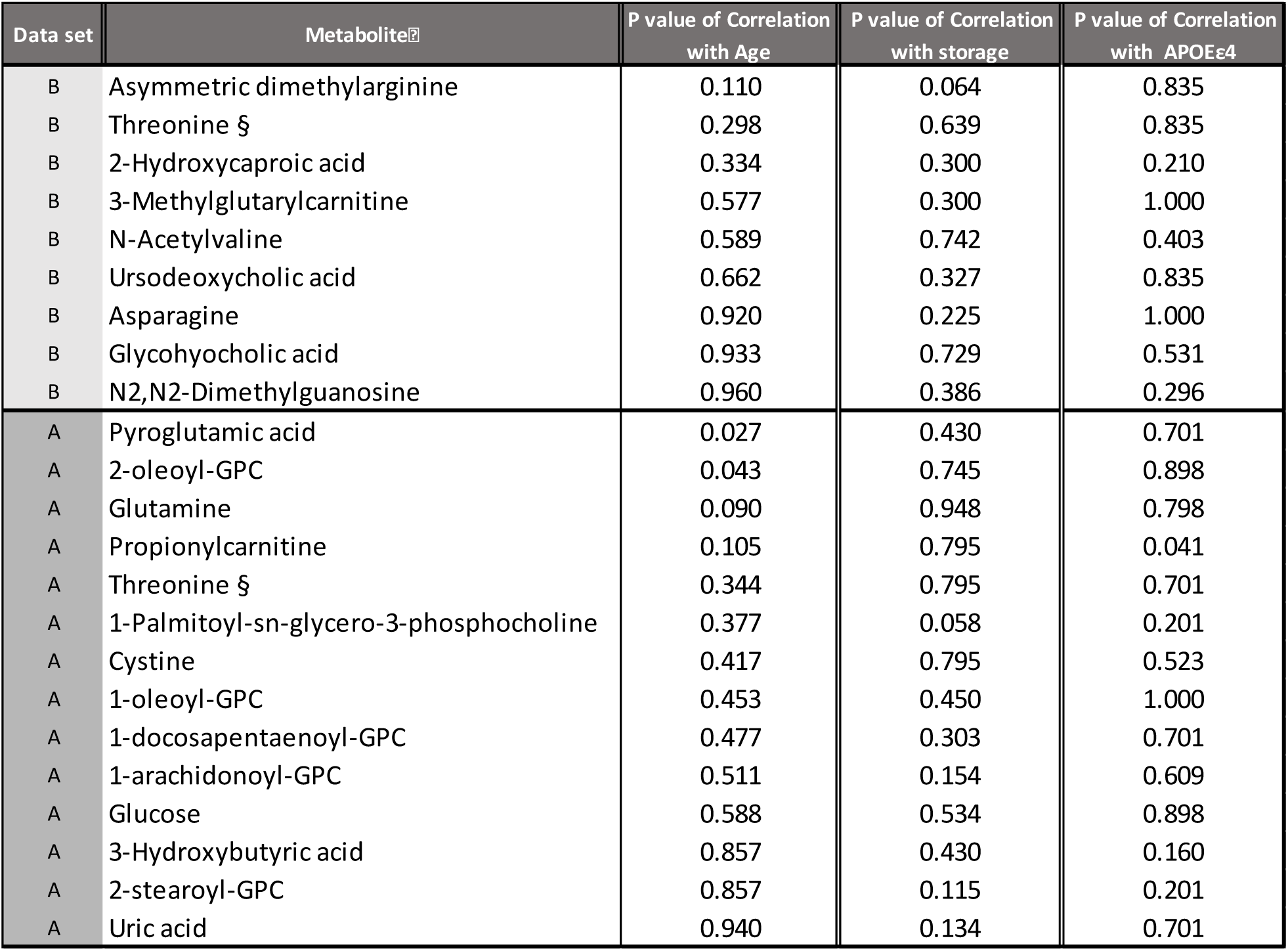
The Spearman’s correlation between 23 metabolites and age at the time of sampling, number of storage years and presence of *APOEε4* allele (p<0.05 value was considered as significant).

**Supplementary Table 3:** Spearman correlation between 1199 detected metabolites, in both dataset ‘A’ and ‘B’, and Aβ_1-42/1-40_-ratio. The table shows the unannotated metabolites, presented as ‘mass@retention time’, that shows a significant correlation (p<0.05).

| Data set | Metabolite | Correlation to $A\beta(1-42/1-40)$ | p.value | Data set | Metabolite | Correlation to $A\beta(1-42/1-40)$ | p.value |
| --- | --- | --- | --- | --- | --- | --- | --- |
| A | 266.0853@0.91 | 0.769 | 0.002 | B | 145.1099@0.4 | 0.867 | 0.001 |
| A | 511.3268@5.98 | -0.764 | 0.002 | B | 398.2297@5.62 | -0.855 | 0.002 |
| A | 614.3656@4.89 | -0.758 | 0.003 | B | 136.0383@0.82 | -0.782 | 0.008 |
| A | 144.0335@0.91 | 0.742 | 0.004 | B | 128.0582@0.39 | 0.770 | 0.009 |
| A | 625.3457@4.9 | -0.709 | 0.007 | B | 256.8292@3.04 | -0.760 | 0.011 |
| A | 310.0491@0.9 | 0.709 | 0.007 | B | 293.1471@1.59 | 0.758 | 0.011 |
| A | 238.1416@2.17 | -0.703 | 0.007 | B | 523.3646@6.82 | -0.758 | 0.011 |
| A | 133.9954@0.58 | -0.698 | 0.008 | B | 143.0947@0.4 | -0.758 | 0.011 |
| A | 247.0843@2.73 | -0.665 | 0.013 | B | 271.9675@0.59 | -0.745 | 0.013 |
| A | 2184.6055@4.97 | 0.659 | 0.014 | B | 480.3801@7.14 | 0.733 | 0.016 |
| A | 168.0284@0.77 | -0.659 | 0.014 | B | 2354.6249@4.48 | -0.732 | 0.016 |
| A | 1266.4967@4.95 | 0.648 | 0.017 | B | 258.8263@3.04 | -0.711 | 0.021 |
| A | 174.0889@3.39 | -0.643 | 0.018 | B | 258.1828@5.28 | -0.709 | 0.022 |
| A | 286.0922@2.73 | -0.643 | 0.018 | B | 159.0678@1.53 | -0.701 | 0.024 |
| A | 312.1756@6.5 | 0.637 | 0.019 | B | 190.0467@1.08 | -0.697 | 0.025 |
| A | 286.2139@5.89 | -0.637 | 0.019 | B | 523.3645@6.73 | -0.697 | 0.025 |
| A | 569.3488@6.21 | 0.637 | 0.019 | B | 204.0092@2.69 | -0.690 | 0.027 |
| A | 257.1029@0.43 | 0.637 | 0.019 | B | 625.3456@4.96 | -0.685 | 0.029 |
| A | 596.2955@6.38 | -0.615 | 0.025 | B | 117.0786@1.63 | 0.685 | 0.029 |
| A | 264.111@2.73 | -0.61 | 0.027 | B | 129.0791@0.59 | -0.685 | 0.029 |
| A | 159.1256@0.4 | 0.61 | 0.027 | B | 482.2873@5.47 | -0.681 | 0.030 |
| A | 567.3535@6.45 | 0.599 | 0.031 | B | 346.1623@3.84 | -0.681 | 0.030 |
| A | 358.0385@0.78 | -0.599 | 0.031 | B | 272.2352@6.82 | -0.673 | 0.033 |
| A | 176.0944@1.26 | -0.598 | 0.031 | B | 463.1833@4.05 | -0.673 | 0.033 |
| A | 312.0614@0.87 | -0.588 | 0.035 | B | 353.0925@4.12 | -0.673 | 0.033 |
| A | 285.194@3.74 | -0.588 | 0.035 | B | 231.1469@1.82 | -0.673 | 0.033 |
| A | 246.056@3.11 | 0.584 | 0.036 | B | 309.1931@3.92 | -0.669 | 0.034 |
| A | 147.9903@0.59 | -0.582 | 0.037 | B | 463.1834@4.05 | -0.661 | 0.038 |
| A | 548.0479@0.78 | -0.582 | 0.037 | B | 187.0634@2.35 | -0.656 | 0.039 |
| A | 145.0524@1.59 | -0.582 | 0.037 | B | 2355.1256@4.48 | -0.656 | 0.039 |
| A | 1100.6119@6.12 | 0.577 | 0.039 | B | 446.2297@5.64 | -0.653 | 0.041 |
| A | 146.0689@2.73 | -0.571 | 0.041 | B | 136.0522@2.2 | -0.648 | 0.043 |
| A | 1284.2694@4.95 | 0.571 | 0.041 | B | 166.0629@3.31 | -0.648 | 0.043 |
| A | 557.3319@5.08 | -0.56 | 0.046 | B | 120.0572@3.31 | -0.648 | 0.043 |
| A | 1284.6321@4.95 | 0.56 | 0.046 | B | 379.2486@5.93 | -0.648 | 0.043 |
| A | 738.0579@0.77 | -0.56 | 0.047 | B | 279.1312@0.94 | 0.636 | 0.048 |
| A | 298.1601@6.09 | 0.555 | 0.049 | B | 246.2347@5.93 | -0.636 | 0.048 |
| A | 521.3489@6.46 | 0.555 | 0.049 |  |  |  |  |

**Supplementary Table 4: (excel file)** List of upstream regulators, which were enriched for the 23 annotated metabolites in the Ingenuity Pathway Analysis (IPA). These regulators include enzymes, kinases, growth factor and transcription factors.

## Supplementary methods

### Metabolite profiling of Plasma

#### Sample preparation

Sample preparation of plasma was performed according to A et al [58]. In detail, 900 µL of extraction buffer (90/10 v/v methanol: water), including internal standards for both GC-MS and LC-MS, were added to 100 µL of plasma. The sample was shaken at 30 Hz for 2 minutes in a mixer mill and proteins were precipitated at +4 °C on ice. The sample was centrifuged at +4 °C, 14 000 rpm, for 10 minutes. The supernatant, 200µL for LC-MS analysis and 50µL to GC-MS analysis, was transferred to micro vials and evaporated to dryness in a speed-vac concentrator. Solvents were evaporated and the samples were stored at −80 °C until analysis.

A small aliquot of the remaining supernatant was pooled and used to create quality control (QC) samples. MSMS analysis (LC-MS) was run on the QC samples for identification purposes. The samples were analyzed in batches according to a randomized run order on both GC-MS and LC-MS.

#### GC-MS analysis

Derivatization and GC-MS analysis were performed as described previously [58]. 0.5 μL of the derivatized sample was injected in splitless mode by a L-PAL3 autosampler (CTC Analytics AG, Switzerland) into an Agilent 7890B gas chromatograph equipped with a 10 m x 0.18 mm fused silica capillary column with a chemically bonded 0.18 μm Rxi-5 Sil MS stationary phase (Restek Corporation, USA) The injector temperature was 270 °C, the purge flow rate was 20 mL min-1 and the purge was turned on after 60 seconds. The gas flow rate through the column was 1 mL min-1, the column temperature was held at 70 °C for 2 minutes, then increased by 40 °C min-1 to 320 °C and held there for 2 minutes. The column effluent was introduced into the ion source of a Pegasus BT time-of-flight mass spectrometer, GC/TOFMS (Leco Corp., St Joseph, MI, USA). The transfer line and the ion source temperatures were 250 °C and 200 °C, respectively. Ions were generated by a 70eV electron beam at an ionization current of 2.0 mA, and 30 spectra s-1 were recorded in the mass range m/z 50 - 800. The acceleration voltage was turned on after a solvent delay of 150 seconds. The detector voltage was 1800-2300 V.

#### LC-MS analysis

Before LC-MS analysis the sample was re-suspended in 10 + 10 µL methanol and water. Each batch of samples was first analyzed in positive mode. After all samples within a batch had been analyzed, the instrument was switched to the negative mode and the second injection of each sample was performed.

The chromatographic separation was performed on an Agilent 1290 Infinity UHPLC-system (Agilent Technologies, Waldbronn, Germany). 2 μL of each sample were injected onto an Acquity UPLC HSS T3, 2.1 x 50 mm, 1.8 μm C18 column in combination with a 2.1 mm x 5 mm, 1.8 μm VanGuard precolumn (Waters Corporation, Milford, MA, USA) held at 40 °C. The gradient elution buffers were A (H2O, 0.1 % formic acid) and B (75/25 acetonitrile:2-propanol, 0.1 % formic acid), and the flow-rate was 0.5 ml min-1. The compounds were eluted with a linear gradient consisting of 0.1 - 10 % B over 2 minutes, B was increased to 99 % over 5 minutes and held at 99 % for 2 minutes; B was decreased to 0.1 % for 0.3 minutes and the flow-rate was increased to 0.8 mL min-1 for 0.5 minutes; these conditions were held for 0.9 minutes, after which the flow-rate was reduced to 0.5 mL min-1 for 0.1 minutes before the next injection.

The compounds were detected with an Agilent 6550 Q-TOF mass spectrometer equipped with a jet stream electrospray ion source operating in positive or negative ion mode. The settings were kept identical between the modes, with the exception of the capillary voltage. A reference interface was connected for accurate mass measurements; the reference ions purine (4 μM) and HP-0921 (Hexakis (1H, 1H, 3H-tetrafluoropropoxy) phosphazine) (1 μM) were infused directly into the MS at a flow rate of 0.05 mL min-1 for internal calibration, and the monitored ions were purine m/z 121.05 and m/z 119.03632; HP-0921 m/z 922.0098 and m/z 966.000725 for positive and negative mode respectively. The gas temperature was set to 150°C, the drying gas flow to 16 L min-1 and the nebulizer pressure 35 psig. The sheath gas temp was set to 350°C and the sheath gas flow 11 L min-1. The capillary voltage was set to 4000 V in positive ion mode, and to 4000 V in negative ion mode. The nozzle voltage was 300 V. The fragmentor voltage was 380 V, the skimmer 45 V and the OCT 1 RF Vpp 750 V. The collision energy was set to 0 V. The m/z range was 70 - 1700, and data was collected in centroid mode with an acquisition rate of 4 scans s-1 (1977 transients/spectrum).

#### Data analysis evaluation/statistical methods

For the GC-MS data, all non-processed MS-files from the metabolic analysis were exported from the ChromaTOF software in NetCDF format to MATLAB R2016a (Mathworks, Natick, MA, USA), where all data pre-treatment procedures, such as baseline correction, chromatogram alignment, data compression and Multivariate Curve Resolution were performed using custom scripts. The extracted mass spectra were identified by comparisons of their retention index and mass spectra with libraries of retention time indices and mass spectra [59]. Mass spectra and retention index comparison was performed using NIST MS 2.0 software. The annotation of mass spectra was based on reverse and forward searches in the library. Masses and ratio between masses indicative of a derivatized metabolite were especially notified. If the mass spectrum according to SMC’s experience was with the highest probability indicative of a metabolite and the retention index between the sample and library for the suggested metabolite was ± 5 (usually less than 3) the deconvoluted “peak” was annotated as an identification of a metabolite.

For the LC-MS data, all data processing was performed using the Agilent Masshunter Profinder version B.08.00 (Agilent Technologies Inc., Santa Clara, CA, USA). The processing was performed both in a target and an untargeted fashion. For target processing, a pre-defined list of metabolites commonly found in plasma and serum were searched for using the Batch Targeted feature extraction in Masshunter Profinder. An in-house LC-MS library built up by authentic standards run on the same system with the same chromatographic and mass-spec settings, were used for the targeted processing. The identification of the metabolites was based on MS, MS-MS and retention time information. For the untargeted data, each group was processed individually using the Batch Recursive Feature Extraction algorithm within Masshunter Profinder.

### Solvents and standards used

#### Solvents

Methanol, HPLC-grade was obtained from Fischer Scientific (Waltham, MA, USA) Chloroform, Suprasolv for GC was obtained from Merck (Darmstadt, Germany) Acetonitrile, HPLC-grade was obtained from Fischer Scientific (Waltham, MA, USA) 2-Propanol, HPLC-grade was obtained from VWR (Radnor, PA, USA) H2O, Milli-Q.

#### Reference and tuning standards

Purine, 4 μM, Agilent Technologies (Santa Clara, CA, USA) HP-0921 (Hexakis(1H, 1H, 3H-tetrafluoropropoxy)phosphazine), 1 μM, Agilent Technologies (Santa Clara, CA, USA) Calibrant, ESI-TOF, ESI-L Low Concentration Tuning Mix, Agilent Technologies (Santa Clara, CA, USA) HP-0321 (Hexamethoxyphosphazine), 0.1 mM, Agilent Technologies (Santa Clara, CA, USA).

#### Stable isotopes internal standards

LC-MS internal standards: 13C9-Phenylalanine, 13C3-Caffeine, D4-Cholic acid, D8-Arachidonic Acid, 13C9-Caffeic Acid were obtained from Sigma (St. Louis, MO, USA). GC-MS internal standards: L-proline-13C5, alpha-ketoglutarate-13C4, myristic acid-13C3, cholesterol-D7 were obtained from Cil (Andover, MA, USA). Succinic acid-D4, salicylic acid-D6, L-glutamic acid-13C5,15N, putrescine-D4, hexadecanoic acid-13C4, D-glucose-13C6, D-sucrose-13C12 were obtained from Sigma (St. Louis, MO, USA).

